# Dystrophin-deficiency stiffens skeletal muscle and impairs elasticity: an *in vivo* rheological examination

**DOI:** 10.1101/2025.05.25.655634

**Authors:** Pavithran Devananthan, Rebecca Craven, Kellie Joe, Gretel S. Major, Jiayi Chen, Natalia Kabaliuk, Angus Lindsay

**Affiliations:** Department of Mechanical Engineering, University of Canterbury, Christchurch, New Zealand; University of Canterbury Biomolecular Interaction Centre, University of Canterbury, Christchurch, New Zealand; School of Biological Sciences, University of Canterbury, Christchurch 8140, New Zealand; Department of Medicine, University of Otago, Christchurch 8014, New Zealand; Maurice Wilkins Centre for Molecular Biodiscovery, Auckland 1010, New Zealand

**Keywords:** Duchenne muscular dystrophy, fibrosis, muscle stiffness, skeletal muscle mechanics, rheology, viscoelasticity

## Abstract

Loss of dystrophin alters the biomechanical properties of skeletal muscle, including stiffness. Stiffness is typically assessed passively in excised muscle, but here we present the development of an *in vivo* rheological method to assess the mechanical properties of the tibialis anterior muscle in anaesthetised wild-type (WT; dystrophin-positive) and *mdx* (dystrophin-deficient) mice using a custom-designed apparatus compatible with an MCR 702 rheometer. To characterise stiffness, compressibility, and elasticity, rheological testing included compressive and shear strain protocols, along with recovery and assessments following contraction-induced strength loss. Relative to WT mice, the tibialis anterior of *mdx* mice were thicker, stiffer, and less compressible. These genotype differences aligned with hydroxyproline content, a marker of fibrosis. Post-deformation recovery was impaired in *mdx* mice under shear strain, and eccentric contraction-induced injury further increased stiffness and energy dissipation in the tibialis anterior of *mdx* mice. This rheological platform maintained the *in vivo* integrity of the tibialis anterior muscle of mice and consistently showed that storage and loss moduli can sensitively detect the detrimental impact of dystrophin-deficiency on the *in vivo* viscoelastic properties of skeletal muscle. This rheological platform, termed myomechanical profiling could be a viable and sensitive tool for assessing muscle quality and mechanical behaviour of skeletal muscle where viscoelastic properties are affected by disease.

## Introduction

Dystrophin is critical for maintaining the structural integrity of muscle fibres [1–3] via its complexation within the dystrophin-glycoprotein complex (DGC). The DGC biomechanically links the actin cytoskeleton to the extracellular matrix [4], stabilizing the sarcolemma, distributing mechanical stress during muscle contraction, and protecting the cell from contraction-induced damage [5, 6]. In healthy muscle, dystrophin acts as a molecular shock absorber, ensuring that muscle fibres maintain their shape and function under repeated cycles of contraction and relaxation [6]. When dystrophin is absent or non-functional due to variants in the Duchenne muscular dystrophy (DMD) gene, muscle fibres experience repeated mechanical damage, leading to cycles of injury and degeneration. Over time, the regenerative capacity of the muscle is overwhelmed, resulting in progressive fibrosis and fat infiltration, which further compromises muscle function [7–9].

The loss of dystrophin has significant consequences for muscle mechanical properties, such as increased stiffness due to excessive and disrupted collagen deposition and architecture [1, 2, 4, 5]. An increase in stiffness alters the viscoelastic behaviour of dystrophin-deficient muscle, reducing its ability to absorb mechanical energy and increasing susceptibility to contraction-induced injury [10]. Fat infiltration further exacerbates these mechanical deficiencies by replacing functional contractile tissue with non-contractile material, weakening the muscle’s force-generating capacity [9]. These biomechanical changes result in reduced elasticity, impaired contraction efficiency, and an overall decline in muscle function, ultimately contributing to the severe physical impairments observed in DMD [11]. Additionally, the absence of dystrophin leads to alterations in key mechanical properties such as increased passive stiffness, reduced compliance, and a lower capacity for energy dissipation during mechanical loading [12]. This manifests as an increase in the storage modulus, indicating a stiffer and less deformable muscle, while the loss modulus may decrease due to reduced ability to dissipate energy. These changes collectively affect muscle resilience, likely leading to heightened susceptibility to mechanical failure.

Typically, the viscoelastic properties of dystrophin-positive and dystrophin-negative skeletal muscle are quantified using excised muscle of the lower hindlimb of mice, where active or passive mechanical properties of dystrophin-negative skeletal muscle are stiffer and less compliant when forces are applied along- or across-muscle fibres in a controlled microenvironment (e.g., Biotester or *ex vivo* chamber with a Krebs buffer) [3, 13–19]. Studies applying dynamic and compressive forces to single fibres or whole muscle using atomic force microscopy or indirect magnetic resonance elastography, have also shown that dystrophin-deficiency compromises the viscoelastic properties of skeletal muscle [20–22]. Passive mechanical properties of the anterior and posterior muscles of the lower hindlimb using rotation about the ankle *in vivo* have also been used as a readout of therapeutic efficacy for a vasomodulator/antioxidant in the *mdx* mouse model of DMD [23]. These techniques typically apply forces in a single plain, which can be challenging to interpret given the various pennation angles of skeletal muscle architecture [24]. Additionally, dystrophin provides a biomechanical link between the intracellular actin cytoskeleton and extracellular matrix and acts like a molecular shock absorber [6, 25], which current techniques might not be able to quantify because there is no assessment of compression. To fully comprehend the effect of dystrophin-deficiency on skeletal muscle mechanics, as well as any injury or exercise-induced exacerbation of the dystrophic microenvironment, a technique to measure the whole muscle response to compression and rotational strain *in vivo* is required.

Rheology is a technique commonly used to characterize the viscoelastic properties of materials, primarily in the context of liquids and soft solids [26]. Rheology quantifies how materials deform and flow under applied stress, providing insight into their mechanical behaviour [27]. While traditionally applied in polymer science and fluid dynamics, rheology has been increasingly utilized in biomedical research to assess the mechanical properties of biological tissues [28–31]. A recent study has demonstrated the potential of rheology for evaluating muscle stiffness and viscoelasticity in a murine model of DMD [22]. Dystrophin-positive skeletal muscle exhibited a higher storage (elastic response) and loss (viscous response) modulus allowing it to effectively absorb and distribute mechanical forces [22]. In contrast, dystrophin-deficient muscle displayed a lower storage and loss modulus, indicating an impaired ability to absorb and dissipate energy [22]. However, this study again used isolated and excised muscle that does not account for the impact of the *in vivo* environment on skeletal muscle viscoelasticity.

In this study, we have developed a customised *in vivo* platform that conforms to the constraints of a live adult mouse to measure the viscoelastic properties of dystrophin-positive and dystrophin-negative skeletal muscle using rheology. The platform can measure the viscoelastic properties of the tibialis anterior (TA) muscle *in vivo* to compressive and rotational strain with micro-newton precision. The customised rheological approach confirms dystrophin-deficiency stiffens skeletal muscle, impairs its elasticity and that contraction-induced damage exacerbates these viscoelastic properties. This approach represents a novel refinement over *ex vivo* mechanical testing methods, offering more physiologically relevant data while minimizing tissue disruption in the context of DMD.

## Methods

### Ethical Approval

The work presented in this study complies with the animal ethics checklist as outlined in Instructions to Authors. All rheological studies were reviewed and approved by the University of Otago Animal Ethics Committee (AUP-24-15). All animals were housed in the Christchurch Animal Research Area (CARA) within the University of Otago, Christchurch (UOC) and treated in accordance with standards set by the University of Otago Animal Welfare Office.

### Experimental mice

All animals were bred in house under a breeding ethics reviewed and approved by the University of Otago Animal Ethics Committee (AUP-23-115). Male C57BL/10ScSn-J (WT) and C57BL/10ScSn-*mdx* (*mdx*) mice between 9 – 14 weeks of age were used for studies. All mice were housed in groups of three to four per cage on a 12/12 h light/dark cycle with food and water provided *ad libitum*. After rheological profiling was completed on a mouse, they were euthanised by cervical dislocation while under anaesthesia.

### Experimental designs

#### General

All studies described below used the isolated tibialis anterior muscle of anaesthetized mice.

#### Study 1

The purpose of study one was to develop a rheological protocol to measure mechanical properties of mouse skeletal muscles *in vivo*. We selected the starting protocols described below using previous *ex vivo* rheology assessments in dystrophin-positive and dystrophin-negative muscle [22] as well as our own preliminary rheology testing on dystrophin-positive and dystrophin-negative cadaver “*in vivo*” tibialis anterior muscle (data not shown). First, rheological profiling was performed at low levels of pre-compression force and shear strain to minimize any damage to the muscle possibly caused by testing itself while maintaining full contact with the surface of the muscle. Rheological tests were conducted with shear strain amplitudes of 0.001 – 0.5% with a minimum allowable by MCR702e rheometer sensitivity. Recognising strain rate dependency of viscoelastic muscle tissue, and to optimise signal to noise level in the measured data and time to complete tests, testing was performed at shear frequencies of 1, 10 and 20 Hz with 0.01 N compression force. This was followed by rotational shear tests with shear strain extended up to 3% and at an increased compression force of 0.1 N (corresponding to less than 20% of muscle compression) in an attempt to compensate probe slippage. For this, frequencies of 0.1, 1 and 10 Hz were selected as 20 Hz was deemed to result in excessive probe slippage and inconsistent measurements. Only WT mice were used for these experiments (n = 3-7/protocol).

#### Study 2

Using the optimised rheological settings identified in study one, study two aimed to determine the impact of dystrophin-deficiency on the *in vivo* rheological profile of skeletal muscle. Twenty-four hours prior to the rheological profiling, mice were injected with Evan’s blue dye (EBD) to assess muscle damage between WT and *mdx* mice, as well as determining if the parameters used in this study caused any myofibre damage. The tibialis anterior muscle of WT and *mdx* mice (n = 5/genotype) were pre-compressed at 0.1 N and an amplitude sweep with shear strains of 0.001 – 5% at 1 Hz was performed. The mice were then maintained under anaesthesia for a further 30 min, before euthanasia and collection of tibialis anterior muscle for EBD analysis via immunofluorescence. Only the left tibialis anterior muscle was used for rheological evaluation per mouse. The tibialis anterior muscle which was not exposed to rheological profiling was used as a non-compressed control for the EBD comparison.

#### Study 3

Study two determined that dystrophin-deficiency affects the rheological profile (stiffness and elasticity) of skeletal muscle when compressed with 0.1 N (∼13-23% of its original thickness), so study three aimed to further probe this genotypic difference by increasing the compression and standardising the change in muscle thickness between WT and *mdx* mice. The tibialis anterior muscle of WT and *mdx* mice (n = 4-8/genotype) were compressed to 40% of muscle thickness and an amplitude shear sweep of 0.001 – 5% at 1 Hz was performed. The probe was then removed from contact with the tibialis anterior muscle and muscle thickness remeasured using the rheometer 5 min later. The tibialis anterior was collected after the remeasurement of muscle thickness for downstream assessment of hydroxyproline content - a measure of collagen density or fibrosis. This assay was used to test the hypothesis that collagen density, as a product of dystrophin-deficiency-induced degeneration, contributes to the increased muscle stiffness and storage modulus measured in *mdx* mice relative to WT mice.

#### Study 4

Under a standardized compression of 40% loss of muscle thickness, study three determined that *mdx* muscle was stiffer at lower shear strains but became WT-like at higher shear strains, indicating dystrophin-deficiency-induced stiffness is overcome with greater rotational strain. To further probe if this relationship was maintained beyond the 5% shear strain, the tibialis anterior muscle of WT and *mdx* mice (n = 4-6/genotype) was compressed 40% of muscle thickness and an amplitude shear sweep of 0.001 – 10% at 1 Hz was performed. Twenty-four hours prior to the rheological profiling, mice were injected with EBD to assess if the parameters used in this study caused any myofibre damage. The probe was then removed from contact with the tibialis anterior muscle and muscle thickness remeasured 5 min later. The mice were maintained under anaesthesia after the protocol finished, before euthanasia and collection of the tibialis anterior muscles for EBD analysis.

#### Study 5

Studies two to four determined that the loss of dystrophin significantly stiffens skeletal muscle at multiple compression depths and shear strains. However, at 0.1 N compression and 5% shear strain, dystrophin-deficiency does not appear to impact storage modulus or “stiffness”. To determine if repeated high shear strains at a similar starting storage modulus would be impacted by dystrophin-deficiency, in study five the tibialis anterior muscle of WT and *mdx* mice (n = 4-8/genotype) was compressed with 0.1 N and then subjected to 100 x 5% shear strain cycles (not an amplitude sweep). The probe was then removed from contact with the tibialis anterior muscle and muscle thickness remeasured 5 min later.

#### Study 6

The rheological profiling of dystrophin-positive and dystrophin-negative skeletal muscle in the previous studies determined that the degenerative microenvironment of *mdx* mice likely increased its stiffness. To further probe this genotypic disparity, study six compressed tibialis anterior muscle of WT and *mdx* mice (n = 4-6/genotype) to 50% of its original thickness (visually determined to be the maximal change in thickness prior to “bubbling” of the muscle around the rheometer probe), which was then maintained for 30 min. Force was continually measured to determine the change in muscle mechanical properties as the tissue “gave” or began to comply with the compression over time. Twenty-four hours prior to the rheological profiling, mice were injected with EBD to assess if the parameters used in this study caused any myofibre damage. The mice were maintained under anaesthesia for a further 30 min after the protocol finished, before euthanasia and collection of the tibialis anterior muscle for EBD analysis.

#### Study 7

Dystrophin-deficient skeletal muscle is highly susceptible to strength loss and injury induced by eccentric contractions [32] - a phenotype so robust, that many pre-clinical therapeutic studies often test their efficacy to protect against strength loss and injury induced by these damaging contractions. To determine the effect of eccentric contraction-induced injury on the rheological profile of WT and *mdx* muscle, WT and *mdx* mice (n = 4-6/genotype) completed a series of 50 electrically-stimulated eccentric contractions of the anterior crural muscles (includes the tibialis anterior) before completing a rheological characterisation immediately afterward (0.1 N compression with an amplitude shear sweep of bba0.001 – 0.5% at 1 Hz).

### Methods

#### Anaesthesia

Mice were initially anaesthetized in an induction chamber using 5% isoflurane and then maintained by the inhalation of 1-3% isoflurane mixed with oxygen at a flow rate of 100 mL·min−1.

#### Exposure of tibialis anterior

The tibialis anterior muscle of anaesthetised mice was exposed and isolated under microscope from the fascia, extensor digitorum longus muscle and tibia – this approach enabled an isolated rheological characterization of the tibialis anterior without interference from surrounding musculature, tendon or bone.

#### Rheological profiling and apparatus

After isolation of the tibialis anterior, the mouse was transferred under anaesthesia to a test apparatus (**Fig. 1A**) attached to a MCR702e MultiDrive Dynamic Material Analyser rheometer (Anton Paar) (**Fig. 1B**). The mouse in the supine position with an anaesthesia nose cone were supported on a flat platform maintained at 37 ± 2⁰C. The temperature was monitored using an infra-red thermometer. A rigid 80 mm x 4 mm x 0.5 mm stainless steel plank was inserted between the tibialis anterior muscle and the extensor digitorum longus muscle/tibia, placed into the slots in the rest holder posts, and securely clamped in position on both ends (**Fig. 1C**). The height of the bed was adjusted to achieve a 90-degree angle between the femur and tibia. The knee was clamped making sure the tibialis anterior aligned centrally with the probe. The foot was immobilised on and taped to a footrest. The apparatus interfaced the rheometer so that the exposed muscle was located directly under a custom 3 mm cylindrical rheometer probe with the plank aligned centrally with the probe.

**Figure 1.**
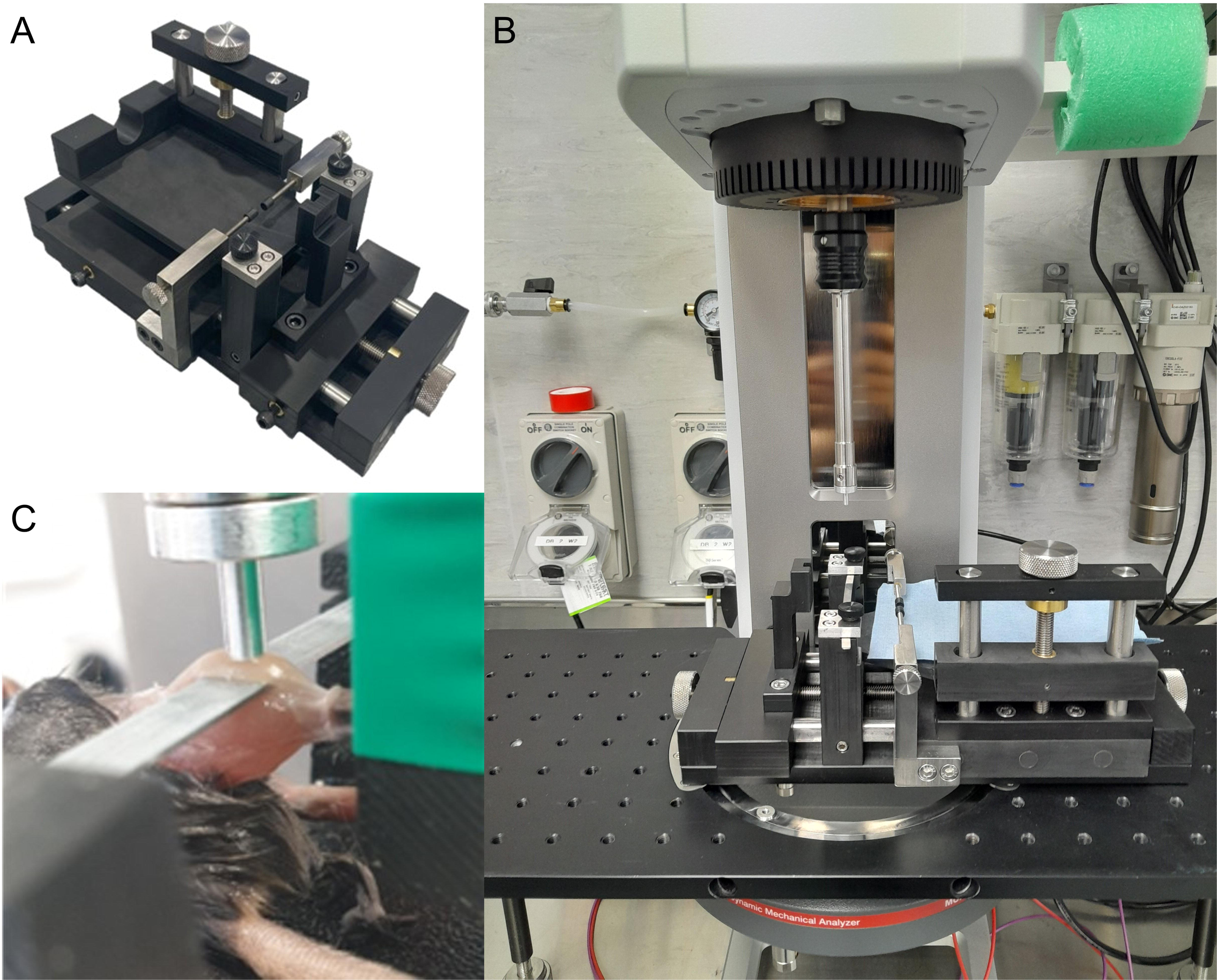
In vivo rheological platform. A) Custom mouse supporting apparatus with different dials to adjust for mouse dimensions (x, y and z planes), B) integrated into an MCR702 rheometer using a threaded board. C) A representative image of a mouse placed onto the apparatus with a plank isolating the tibialis anterior muscle and a probe lowered onto the muscle surface.

In most tests, the tibialis anterior muscle was first pre-compressed by an application of the probe at 10 μm/s to a specified normal force to assure full contact of the probe with the muscle surface. The compressive force was maintained during subsequent dynamic rotational shear sweeping over a range of ascending shear strain amplitudes starting at a shear strain of 0.001% and ramping logarithmically to a maximum of 0.5-10%. The shear strain within the muscle was induced by the probe’s back and forth rotation relative to its original position at a specified frequency. The muscle resistance to this deformation was measured via the storage and loss moduli.

Later targeted tests focused on applying and maintaining a specified compressive strain. Muscle thickness relative to the plank position was measured before, during, and after rheological testing by lowering the probe at a velocity of 10 μm/s until a normal force of 0.01 N was sensed.

#### Muscle damage

EBD was diluted in PBS to 5 mg/mL, filter sterilized with a 0.2 μm filter and injected intraperitoneally at 100 μL/10 g body mass 24 h before the physiological or rheological profiling, as previously described [33]. Tibialis anterior muscles were removed and cryopreserved in liquid nitrogen-cooled isopentane at their respective time-points for each study (described in the experimental designs above).

#### Muscle histology

At the time of sectioning, tibialis anterior muscles were thawed to −20°C and 10 µm mid-belly sections cut. Sections were fixed in −20 °C acetone for 5 min, washed in PBS, blocked for 30 min at room temperature with 5% BSA/PBS, and counterstained with laminin (1:500; Sigma-Aldrich L9393) for 2 h at RT as previously described [34]. Sections were then washed in PBS and incubated with anti-Rabbit Alexa Flour 488 (1:500; ThermoFisher Scientific) for 1 h at RT. Sections were washed a final time in PBS and mounted in ProLong Golf Antifade with DAPI (ThermoFisher Scientific). Images were acquired on a Zeiss AxioImager Z1 fluorescence microscope and stitched together with Zen software (Zeiss) to allow visualization of the entire tibialis anterior. SMASH software was used to determine the percentage of EBD-positive fibres in whole tibialis anterior images as well as quantifying myofibre cross-sectional area and minimum and maximum feret’s diameter [35].

#### Collagen density

Hydroxyproline content of isolated tibialis anterior muscle was measured according to the manufacturer’s instructions (AB222941, Abcam). Muscles were carefully excised to ensure no distal or proximal tendon was left visible that could artificially inflate hydroxyproline content.

#### Muscle physiology

Mice were anaesthetised with isoflurane and maximal isometric torque of the anterior crural muscles was measured as previously described [36, 37] The anaesthetized mouse was placed on a temperature-controlled platform to maintain core temperature at 37°C. The left knee was clamped and the left foot was secured to an aluminium footplate attached to the shaft of the servomotor system (Model 300B-LR; Aurora Scientific, Aurora, Ontario, Canada). Sterilized platinum needle electrodes percutaneously stimulated the left common peroneal nerve, which was connected to the stimulator and stimulus isolation unit. The contractile function of the anterior crural muscles (including the tibialis anterior) was assessed by measuring isometric torque (150 Hz; 150-ms train with 0.1-ms pulses). One minute later, the anterior crural muscles were injured by performing 50 electrically stimulated maximal eccentric contractions, where the foot was passively moved from 0° (positioned perpendicular to tibia) to 19° of dorsiflexion. Then, the anterior crural muscles performed a 100-ms isometric contraction followed by an additional 20 ms of stimulation while the foot was moved to 19° of plantarflexion at 2,000°·s−1. A one min rest after the eccentric contraction protocol was given before reassessing peak isometric torque, then the mouse was transferred under anaesthesia to the rheological setup for profiling as described above.

#### Statistics

Data were analysed in GraphPad Prism 10.0. Data were assessed for normality by the Shapiro-Wilk normality test. Genotype comparisons for a single measure (e.g., muscle thickness) were assessed using an unpaired t-test. For all other data, and if normally distributed, a two- or three-way non-repeated or repeated measures ANOVA or non-parametric equivalent (e.g., Friedman’s test) were used with Bonferroni post-hoc analysis. Linear regression analysis was used to assess the association between force, hydroxyproline content and rheological profiling. All data are presented as mean ± SEM and p < 0.05 was required for significance.

## Results

### Study 1

To optimise a rheological protocol to assess the impact of dystrophin deficiency on muscle stiffness and elasticity *in vivo*, we first designed and engineered an apparatus to support an anaesthetised mouse and exposed tibialis anterior muscle during testing (**Fig. 1A**). The apparatus consisted of a heated bed to rest the body of the mouse and the nose cone and provide body temperature regulation. The bed could be adjusted vertically to accommodate varying femur lengths so that the exposed tibialis anterior was in line with the rheometer probe. The tibialis anterior was supported by a plank inserted underneath the muscle and securely held in place during testing. The plank was manufactured of stainless steel and dimensioned to fit between the tibialis anterior and the extensor digitorum longus muscle without straining or damaging the muscle while being rigid to provide consistent datum for muscle thickness measurements (**Fig. 1C**). The spacing between the plank rest poles was sized to make room for the mouse hindlimb while eliminating the plank flex for the compression forces applied. A footrest and a knee clamp were also incorporated to limit any movement of the mouse during testing. Their position could be adjusted horizontally for varying tibia lengths to maintain ankle and knee joints at 90-degree angles. The knee clamp and the bed were placed onto the same steel lead screw rail slider system. The knee clamp was made from centred M3 threaded steel rods and supported on the L-shaped holders. The footrest was designed to the average dimensions of a mouse foot. The apparatus was made from black acetal, with the parts undergoing rotation made from steel and brass. The test apparatus, capable of stabilising an isolated tibialis anterior muscle of an adult mouse while under anaesthesia, was fitted to the rheometer (**Fig. 1B**) and multiple testing protocols were assessed to elucidate the rheological profile of a skeletal muscle.

Under 0.01 N compressive force and an amplitude shear strain sweep between 0.001 – 0.5% at 1 – 20 Hz, there was a main effect of frequency on storage modulus (p = 0.021) but no effect of shear strain (**Fig. 2A**; p = 0.621). A frequency of 20 Hz resulted in greater storage modulus across the shear strain amplitude relative to 1 and 10 Hz (**area under the curve [AUC];** p < 0.001), but the higher frequencies also resulted in larger inter-mouse variation, likely due to lower signal to noise levels associated with periodic slippage. Loss modulus showed a similar trend to storage modulus, with a frequency main effect approaching significance (**Fig. 2B**; p = 0.065). Loss modulus at 20 Hz was greater across the amplitude shear strain sweep relative to 1 and 10 Hz (**AUC**; p ≤ 0.011), but the 20 Hz frequencies caused repeated slippage of the probe. The loss factor, or the measure of the amount of energy dissipated by the muscle relative to the energy stored elastically when the probe rotates, was also affected by frequency across the amplitude sweep (**Fig. 2C**; p = 0.047), with 20 Hz resulting in a larger loss factor relative to 1 Hz (**AUC**; p = 0.004). Again, the higher frequency-induced slippage of the probe on the muscle generated extreme variation. These data indicate that higher frequencies with 0.01 N compressive force provide inconsistent measurements of muscle mechanics, but to ensure this was not due to the compressive force, we next tested these properties with 0.1 N compressive force, a shear strain between 0.001 – 3% and frequencies between 0.1 – 10 Hz.

**Figure 2.**
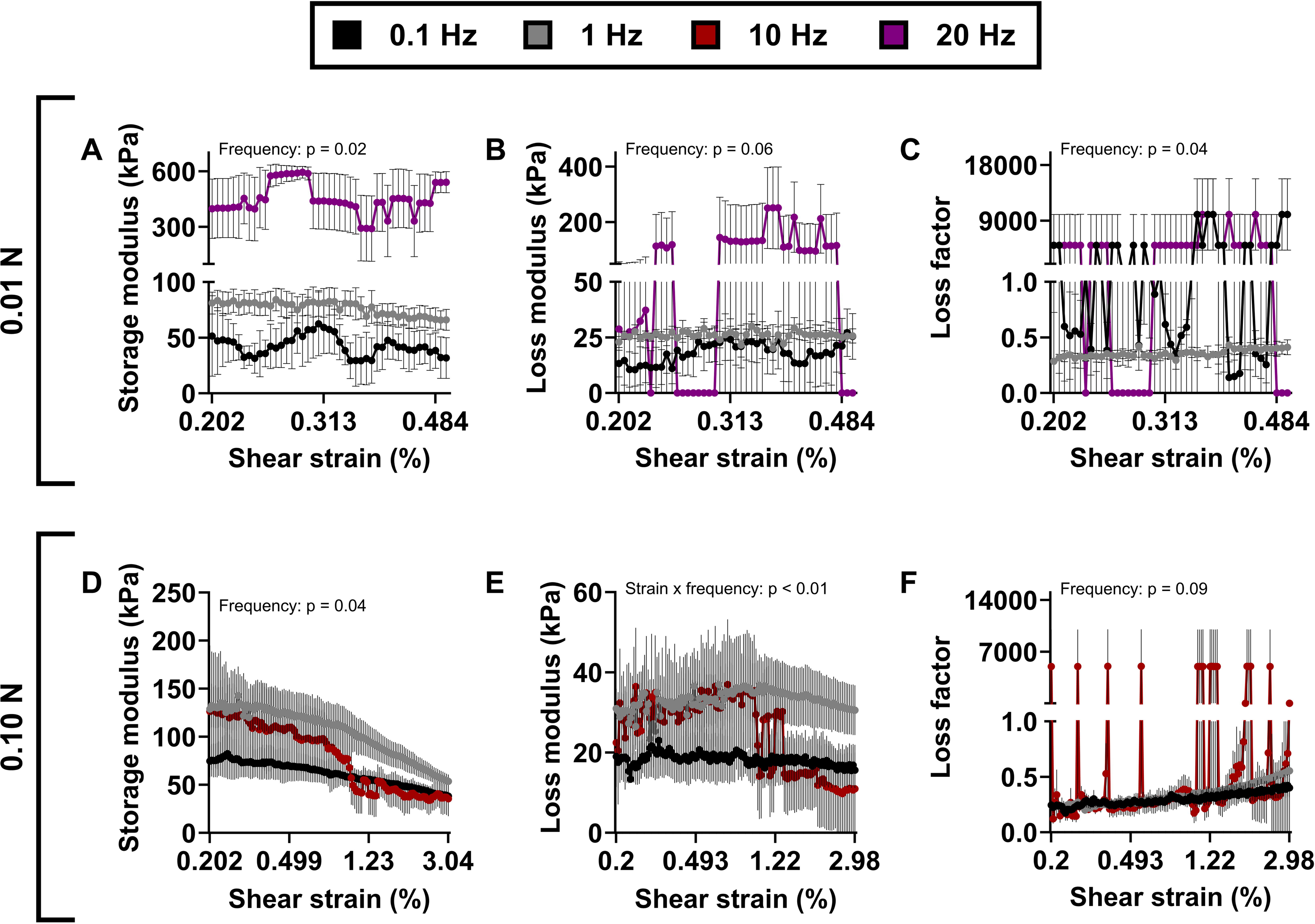
Optimisation of rheological parameters. A) G’ vs shear strain, B) G” vs shear strain and C) loss factor vs shear strain with sweeps between 0.001% - 0.5% and angular shear strain at 0.01N compression. D) G’ vs shear strain, E) G” vs shear strain and F) loss factor vs shear strain with sweeps between 0.001% - 3% angular shear strain at 0.1N compression. Data is mean ± SEM. N = 3-7/protocol.

Relative to 0.01 N compressive force, storage and loss modulus at 1 and 10 Hz were 4-6-fold higher with a 0.1 N compressive force. There was a frequency main effect for storage modulus (**Fig. 2D**; p = 0.045), with the highest values measured for 1 Hz (**AUC**; p ≤ 0.001). Loss modulus was affected by a strain x frequency interaction (**Fig. 2E**; p < 0.001), with values highest for 1 Hz relative to 0.1 and 10 Hz (**AUC**; p ≤ 0.003). At 10 Hz, both storage and loss modulus were affected by slippage of the probe against the muscle at multiple shear strains, resulting in large variation in output – this also resulted in greater loss factor relative to 0.1 and 1 Hz **(Fig. 2F**; p ≤ 0.031).

Collectively, these data show that independent of compressive force, lower frequencies maintain contact between the probe and the muscle, ensuring accurate and consistent readings across increasing shear strains. There is also no difference in rheology performance between 0.1 and 1 Hz, except that output for storage and loss modulus are consistently higher, without slippage. Therefore, to ensure we had the greatest opportunity to distinguish between the rheological characteristics of dystrophin-positive and dystrophin-negative muscle, we selected 0.1 N and 1 Hz parameters across shear strains up to 5% for the following studies.

### Study 2

The optimised protocol (0.1 N, 1 Hz and 0.001 – 5% shear strain) was used to determine differences between WT and *mdx* mice. Tibialis anterior of *mdx* mice was thicker relative to WT mice (**Fig. 3A**; p < 0.001) and was compressed less under 0.1 N (**Fig. 3B**; muscle thickness x genotype: p < 0.001). There was a shear strain x genotype interaction for storage modulus (**Fig. 3C**; p < 0.001), with *mdx* tibialis anterior muscle having a greater AUC relative to WT tibialis anterior muscle (**Fig. 3D**; p < 0.001) and storage modulus decreasing with increasing shear strain. The loss modulus was somewhat affected by strain (**Fig. 3E**; p = 0.055) and genotype (p = 0.066), with *mdx* tibialis anterior muscle having a greater AUC relative to WT tibialis anterior muscle (**Fig. 3F**; p < 0.001). Similarly, the loss factor was affected by a shear strain x genotype interaction (**Fig. 3G**; p < 0.001), which increased for both genotypes as shear strain increased, and was greater for *mdx* tibialis anterior muscle (**Fig. 3H**; p < 0.001). At the conclusion of shear strain and a standardised 0.1 N force, *mdx* tibialis anterior had reduced less in thickness than WT tibialis anterior (**Fig. 3I**; p = 0.002). These data indicate *mdx* tibialis anterior is stiffer than WT tibialis anterior, so we next probed this genotypic difference with a greater force that standardised thickness change.

**Figure 3.**
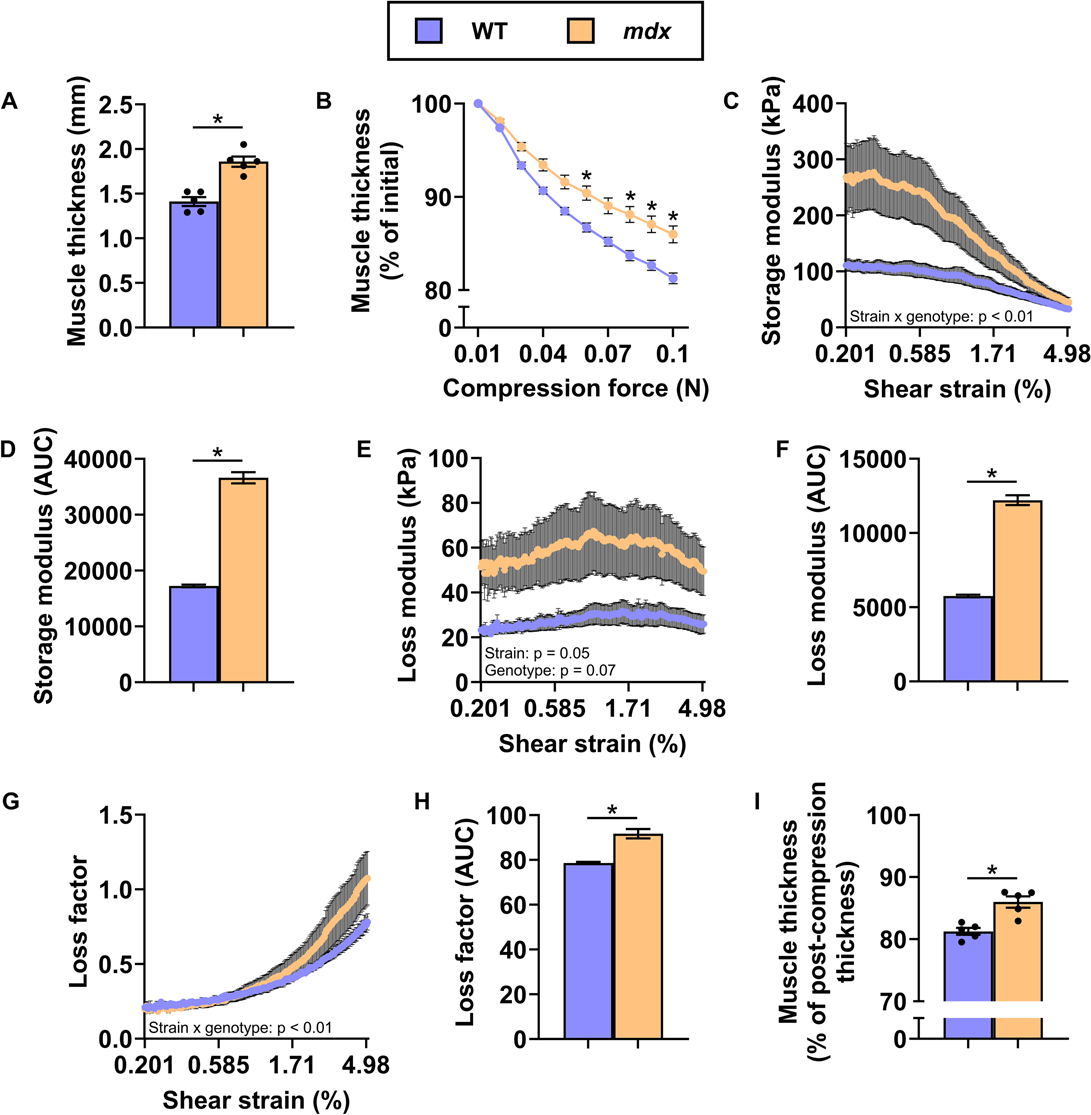
Wildtype (WT) and mdx specimens compressed to 0.1 N with amplitude sweeps between 0.001% and 5% at 1 Hz. A) Initial muscle thickness, B) muscle thickness as a function of compression force, C) storage modulus over amplitude sweep, D) area under the curve of storage modulus, E) loss modulus over amplitude sweep, F) area under the curve of loss modulus, G) loss factor over amplitude sweep, H) area under the curve of loss factor and I) muscle thickness at the end of the test. Data is mean ± SEM. N = 5/genotype. * P < 0.05.

### Study 3

In study 3, we aimed to determine the rheological properties of the tibialis anterior of WT and *mdx* mice to a standardised 40% compression of muscle thickness. The tibialis anterior of *mdx* mice was thicker relative to WT mice (**Fig. 4A**; p < 0.001) and required 54% more force (1.03 vs 0.67 N) to compress 40% of original thickness (**Fig. 4B**; p = 0.002). The storage and loss modulus of the tibialis anterior of WT and *mdx* mice was affected by the greater compressive force relative to 0.1 N in study 2 (∼2-fold increase). However, the same relationship existed between genotypes relative to study 2, with *mdx* tibialis anterior having greater storage (**Fig. 4C, D**; p < 0.001) and loss moduli (**Fig. 4E, F**; p ≤ 0.031) and storage modulus decreased with increasing shear strain (strain x genotype: p < 0.001). The loss factor was not affected by genotype (**Fig. 4G, H**; p ≤ 0.074), but it did increase with shear strain (p < 0.001). Five minutes post-compression, the thickness of the tibialis anterior of WT mice recovered to 90% relative to 82% of *mdx* mice (**Fig. 4I**; p = 0.010). These data indicate that *mdx* muscle is stiffer and it loses elasticity when compressed to a standardised 40% of original thickness relative to WT muscle.

**Figure 4.**
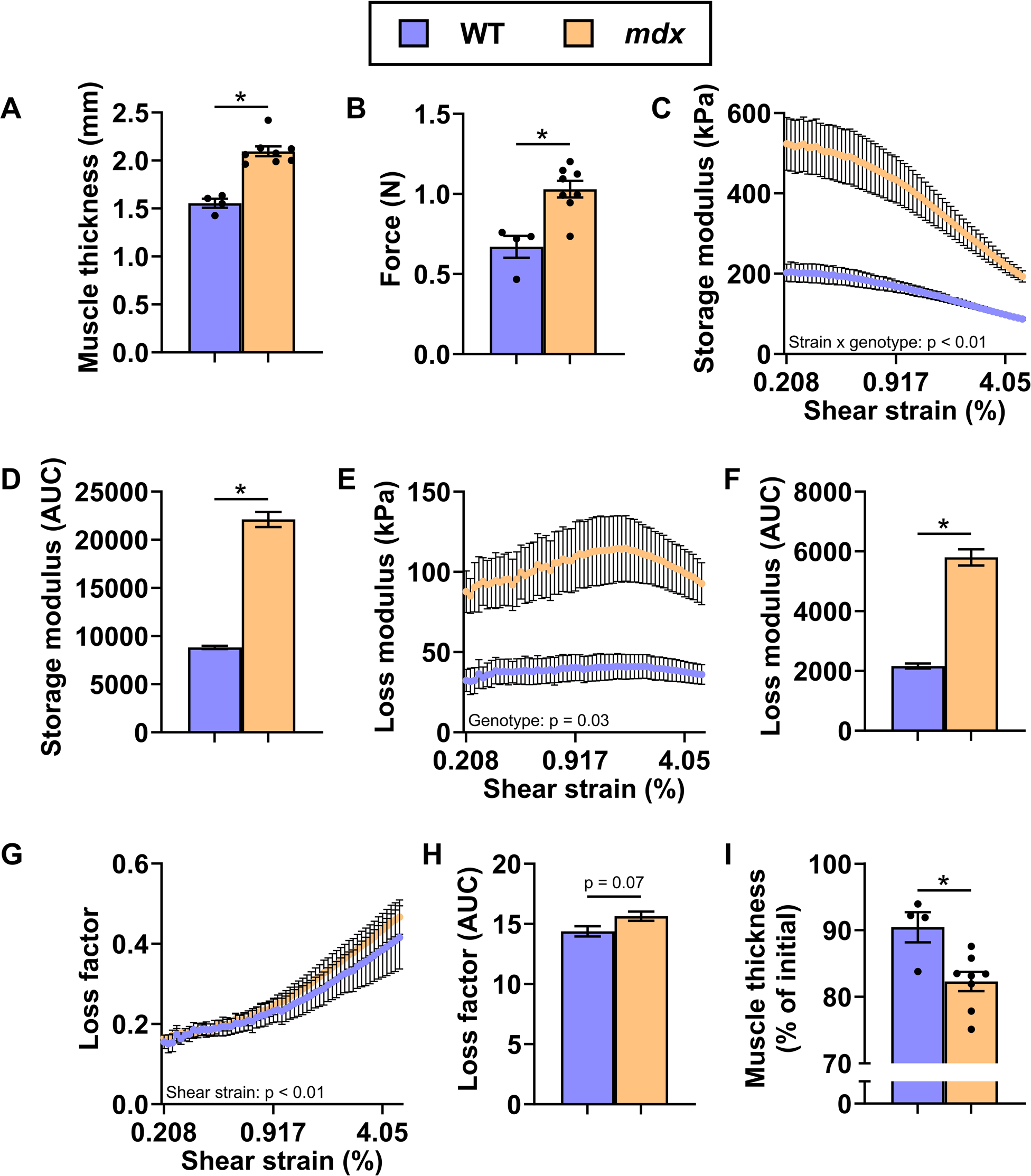
Wildtype (WT) and mdx specimens compressed to 40% of initial muscle thickness, with amplitude sweeps between 0.001% and 5% at 1 Hz. A) Initial muscle thickness, B) compression force to achieve 40% compression, C) storage modulus over amplitude sweep, D) area under the curve of storage modulus, E) loss modulus over amplitude sweep, F) area under the curve of loss modulus, G) loss factor over amplitude sweep, H) area under the curve of loss factor and I) muscle thickness at the end of the test. Data is mean ± SEM. N = 4-8/genotype. * P < 0.05.

To determine the association between the force required to compress the tibialis anterior muscle 40% of original thickness and the muscles rotational stiffness, we completed linear regression analyses for each of the muscles tested in this study as well as muscles tested using the same parameters in Study 4 (up to 5% shear strain only to increase sample size). At 0.2% shear strain, storage and loss modulus were associated with force, but loss factor was not (**Fig. 5A**). At 5% shear strain, the association between storage modulus and force became stronger, loss modulus remained equally associated with force, and loss factor was still not affected (**Fig. 5B**).

**Figure 5.**
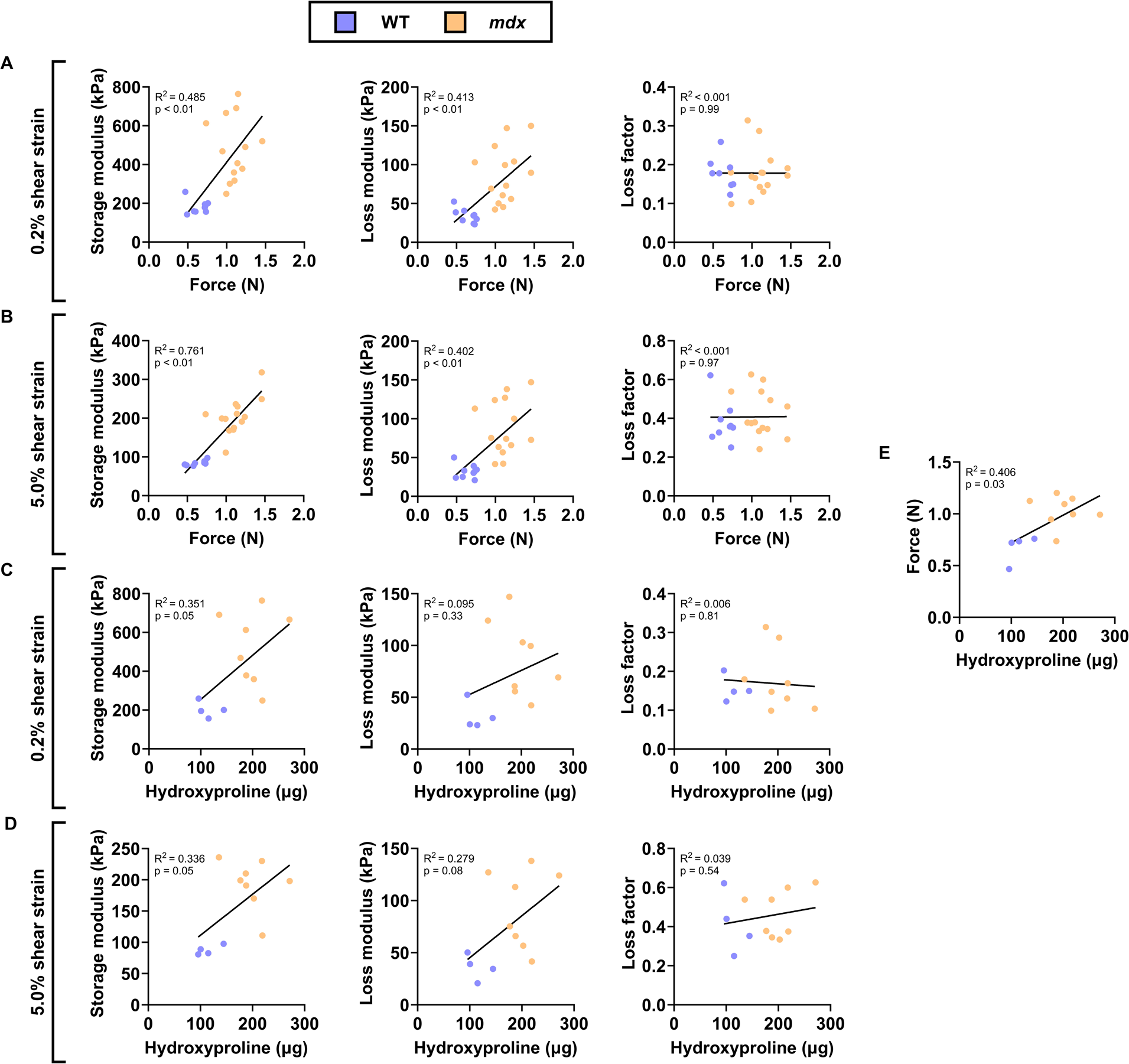
Association between compressive and rotational stiffness and collagen content of skeletal muscle. The association between A) force and G’, G”, and loss factor at 0.2% shear strain, B) force and G’, G”, and loss factor at 5% shear strain, C) hydroxyproline content and G’, G”, and loss factor at 0.2% shear strain, D) hydroxyproline content and G’, G”, and loss factor at 5% shear strain, and E) force and hydroxyproline content of wildtype (WT) and mdx mice.

Because collagen content is a primary pathological hallmark of dystrophin deficiency and the impact of collagen density and architecture (cross-linking) on dystrophic muscle passive stiffness has been heavily investigated and appears to be muscle type dependent [14–17], we determined if the rheological stiffness profile of muscle was influenced by collagen density in muscles from Study 3 (indirectly with hydroxyproline content – absolute hydroxyproline levels are presented because rheology is assessing only a 3 mm diameter section of the mid-belly of the muscle rather than the whole muscle where mass and/or CSA would be important normalisation strategies). *Mdx* tibialis anterior had more hydroxyproline content than WT tibialis anterior (p < 0.001), and the storage modulus was associated with hydroxyproline content at 0.2 and 5% shear strain (**Fig. 5C, D**). Loss modulus and loss factor were not associated with hydroxyproline content (**Fig. 5C, D**). These data suggest that collagen density impacts the stiffness, or resistance to rotational shear using rheology, of the tibialis anterior muscle. Hydroxyproline content was also associated with the force required to compress the tibialis anterior 40% of original thickness (**Fig. 5E**), indicating collagen density also impacts compressive strength of skeletal muscle.

### Study 4

In this study, we maintained the 40% compression parameters but explored the addition of a 10% shear strain. The tibialis anterior muscle of *mdx* mice was thicker (**Fig. 6A**; p < 0.001) and required 2-fold greater force to compress to 40% of original thickness relative to WT mice (**Fig. 6B**; p < 0.001). The storage modulus was ∼2.5-fold greater in *mdx* mice relative to WT mice and decreased with increasing shear strain, which was affected by a shear strain x genotype interaction (**Fig. 6C, D**; p < 0.001). Loss modulus was not affected by shear strain, but it was ∼3-fold greater in *mdx* mice relative to WT mice (**Fig. 6E, F**; p < 0.001). The loss factor increased with shear strain (p < 0.001), which was not different between genotypes across the shear strain amplitude (**Fig. 6G**; p = 0.638), but the AUC showed that *mdx* mice had greater loss factor (**Fig. 6H**; p = 0.006). Muscle thickness 5 min following the removal of the rheometer probe, was not different between genotypes (**Fig. 6I**; p = 0.149). These data indicate that at increased rotational shear, dystrophin-deficient muscle retains its stiffness and capacity to remain elastic.

**Figure 6.**
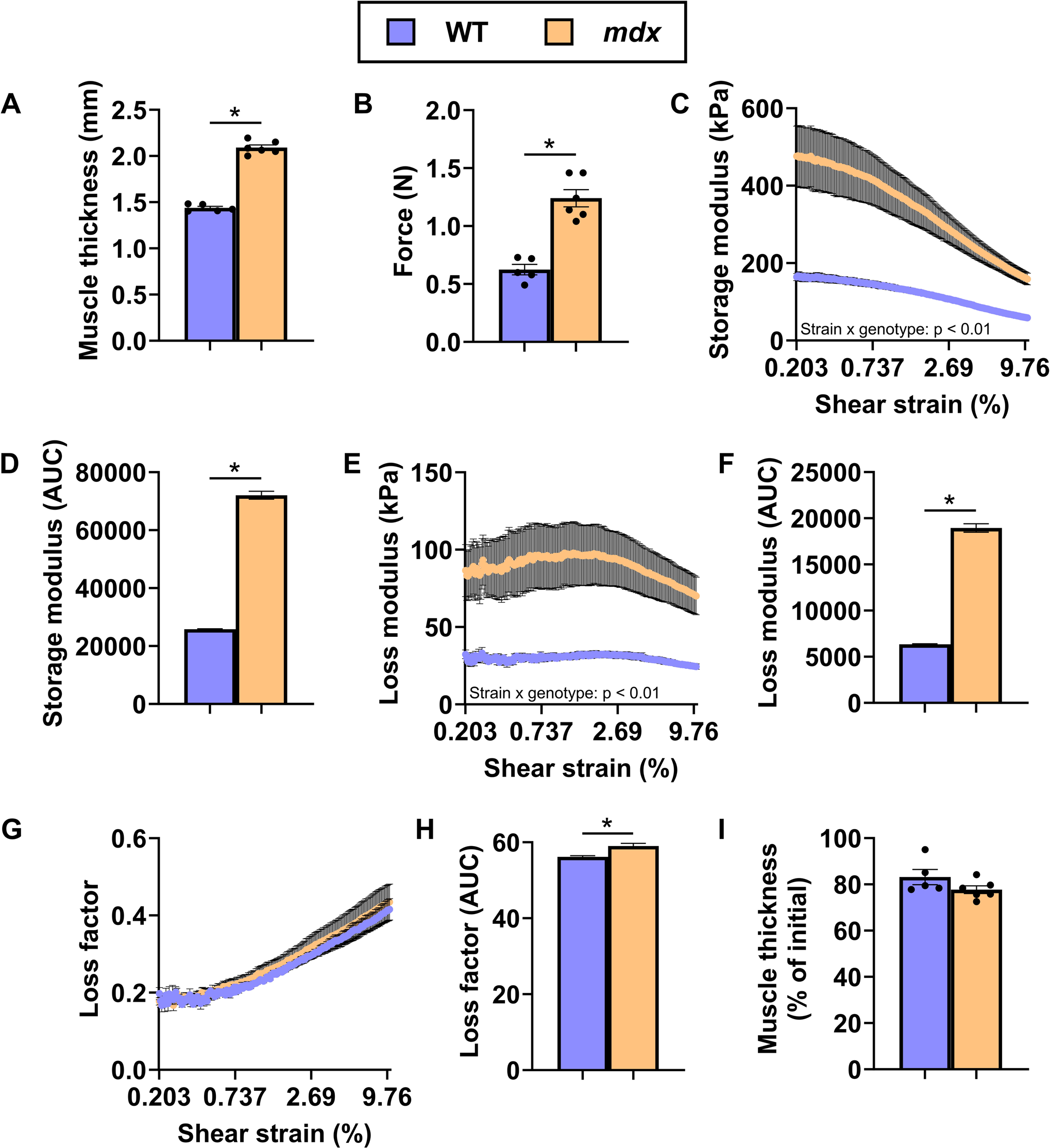
Wildtype (WT) and mdx tibialis anterior compressed to 40% of initial muscle thickness, with amplitude sweeps between 0.001% and 10% at 1 Hz. A) Initial muscle thickness, B) compression force to achieve 40% compression, C) storage modulus over amplitude sweep, D) area under the curve of storage modulus, E) loss modulus over amplitude sweep, F) area under the curve of loss modulus, G) loss factor over amplitude sweep, H) area under the curve of loss factor and I) muscle thickness at the end of the test. Data is mean ± SEM. N = 4-6/genotype. * P < 0.05.

### Study 5

Study 5 aimed to determine if repeated high shear strains at a similar starting storage modulus would be impacted by dystrophin-deficiency. Like study two, *mdx* tibialis anterior muscle in this cohort was thicker (**Fig. 7A**; p < 0.001) and compressed less under 0.1 N force relative to WT (**Fig. 7B**; p = 0.001). At the start of the 100 x 5% shear strain cycles, storage modulus was not different between WT and *mdx* mice, nor did it change (**Fig. 7C, D**; p ≥ 0.716). The loss modulus and loss factor did not change over shear strain cycles for both genotypes (**Fig. 7E, G**; p ≥ 0.119), but these were greater in *mdx* tibialis anterior relative to WT tibialis anterior (**Fig. 7F, H**; p < 0.001). Five minutes post-compression, the thickness of the tibialis anterior of WT mice recovered to 94% relative to 97% of *mdx* mice (**Fig. 7I**; p = 0.013). These data indicate, that even under repeated high shear strains, dystrophin-deficient muscle is not mechanically affected relative to dystrophin-positive muscle.

**Figure 7.**
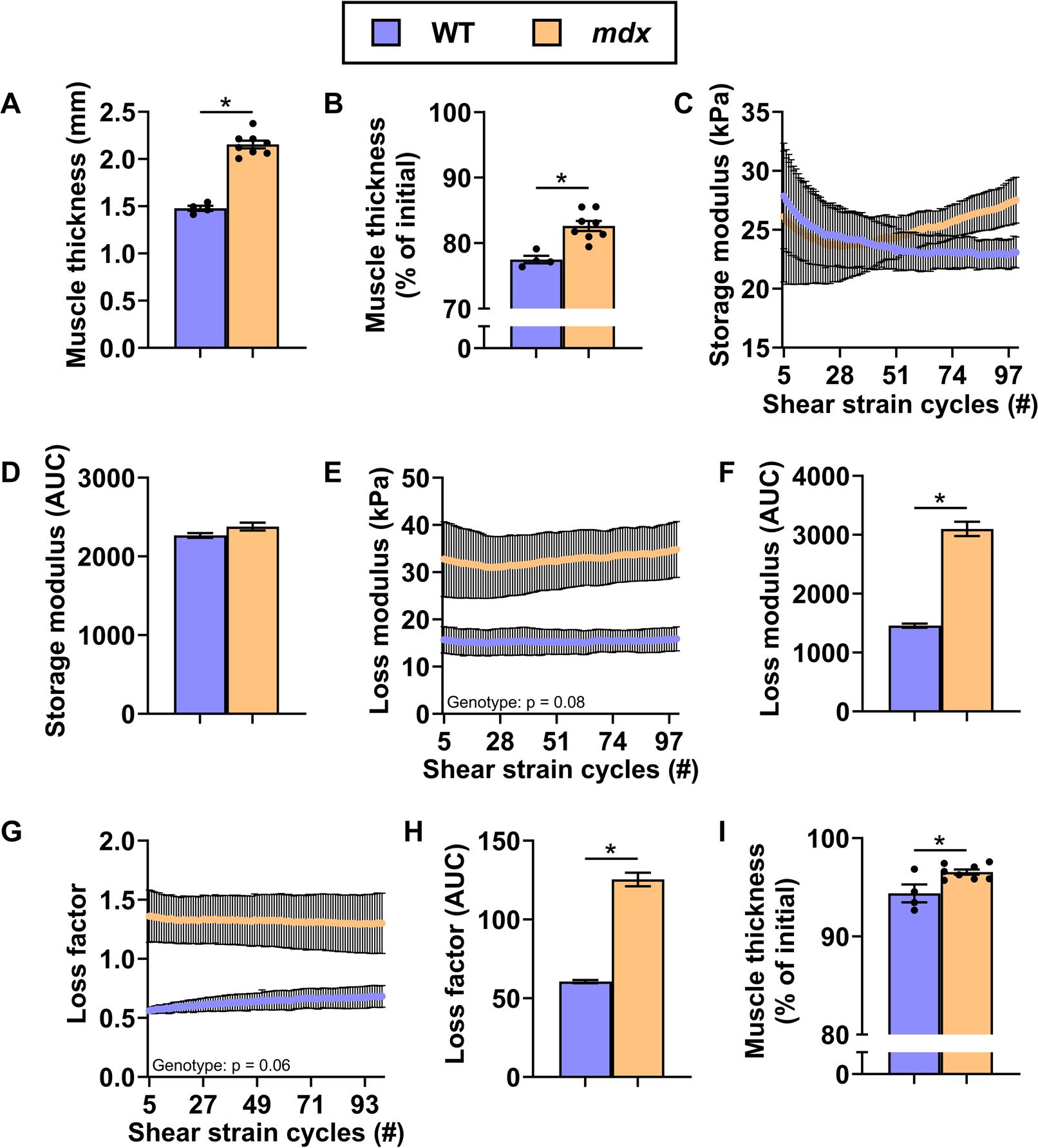
Wildtype (WT) and mdx specimens compressed with 0.1 N, with 100 cycles of 5% shear strain at 1 Hz. A) Initial muscle thickness, B) muscle thickness after 0.1 N compressive force, C) storage modulus over strain cycles, D) area under the curve of storage modulus, E) loss modulus over strain cycles, F) area under the curve of loss modulus, G) loss factor over strain cycles, H) area under the curve of loss factor and I) muscle thickness at the end of test. Data is mean ± SEM. N = 4-8/genotype. * P < 0.05.

### Study 6

This study aimed to determine the impact of dystrophin-deficiency on the relax-ability of muscle to a constant 50% compression strain. Tibialis anterior muscle of *mdx* mice was thicker (**Fig. 8A**; p < 0.001) and required 1.54-fold greater force to compress to 50% of original thickness relative to WT mice (**Fig. 8B**; p < 0.001). Force decreased for both genotypes over time when the muscle thickness was maintained at 50% of its original thickness (**Fig. 8C**), with *mdx* muscle maintaining a higher level of force throughout relative to WT muscle (p < 0.001). However, the rate of change over 30 min, which was affected by a time x genotype interaction (p < 0.001) was not different between genotypes (**Fig. 8D**; p = 0.071). Five minutes following the removal of the rheometer probe after the 30 min test protocol, WT and *mdx* muscle were at 62% and 71% of initial thickness, respectively (**Fig. 8E**; p = 0.003). These data indicate that *mdx* muscle is stiffer, capable of maintaining greater elasticity under high compressive force, but that its capacity to relax under constant high tension is not different than WT mice.

**Figure 8.**
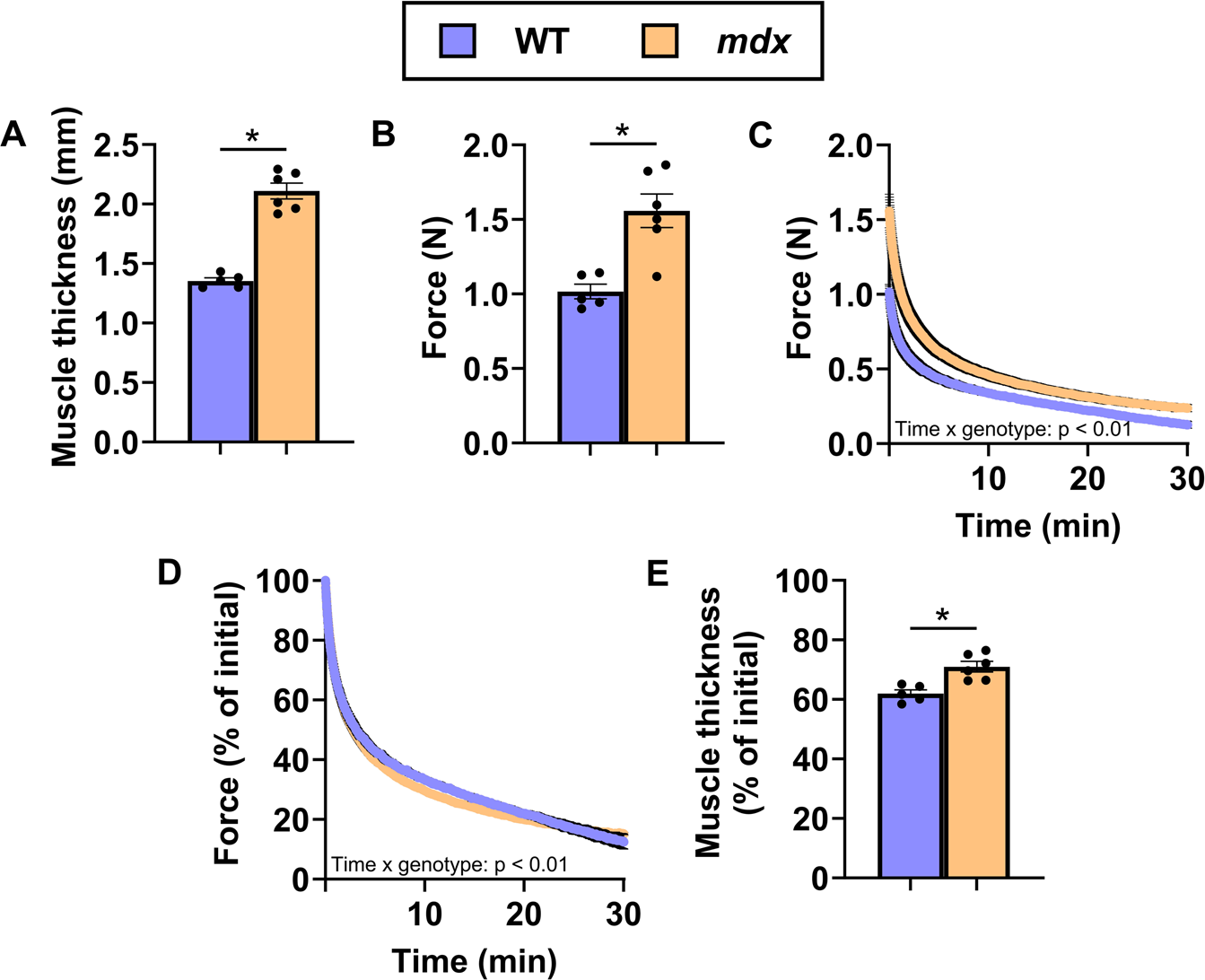
Wildtype (WT) and mdx specimens compressed to 50% of initial muscle thickness and left compressed for 30 min. A) Initial muscle thickness, B) compression force to achieve 50% compression, C) normal force reading from muscle over time, D) percent of initial force over time and E) muscle thickness at the end of the test. Data is mean ± SEM. N = 4-6/genotype. * P < 0.05.

### Muscle damage and muscle fibre morphology

Under control conditions, 1.25% of *mdx* muscle fibres were positive for EBD relative to 0% of WT muscle fibres (**Fig. 9A, B**). The number of EBD positive fibres for both genotypes did not change when compressed 0.1 N with 5% shear strain (p = 0.999). When compressed 40% or 50% with 10% shear strain or for 30 min respectively, 20% of muscle fibres from both genotypes were positive for EBD on the periphery and in the centre of the cross section, with an overall effect of compression (**Fig. 9A, B**; p < 0.001).

**Figure 9.**
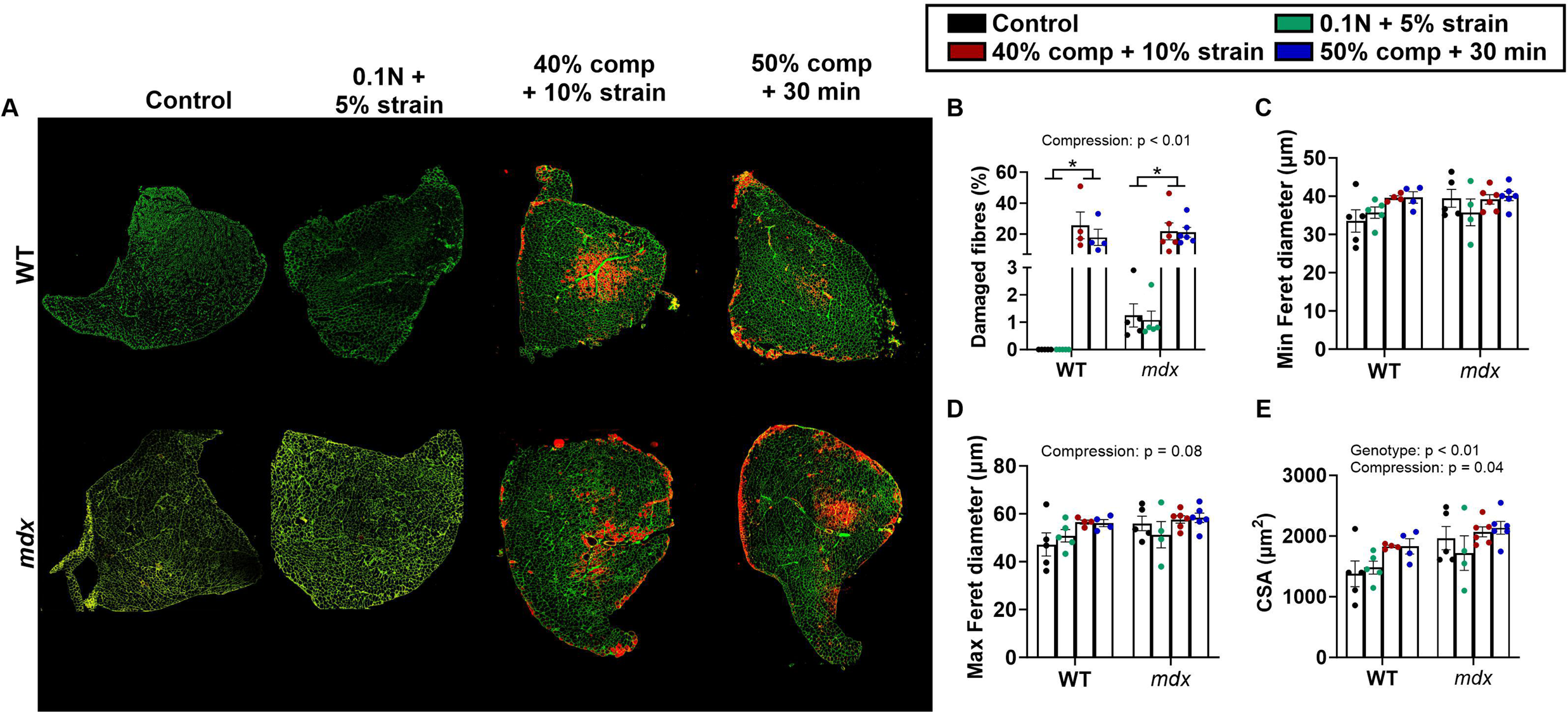
High rheological compression damages skeletal muscle. A) Representative images of wildtype (WT) and mdx Evan’s Blue Dye-stained skeletal muscle cross-sections under various rheological parameters. B) Percent of damaged muscle fibres. C) Minimum Feret diameter of muscle fibres. D) Maximum Feret diameter of muscle fibres. E) Cross sectional area of muscle fibres.

Minimum and maximum Feret’s diameter of tibialis anterior muscle fibres was not different between genotypes (p ≥ 0.119), nor did the type of rheological protocol have an effect (**Fig. 9A, C, D**; p ≥ 0.083). Under control conditions, the average CSA of the tibialis anterior muscle fibres was greater in *mdx* mice relative to WT mice (1964 vs 1376 µm^2^). After rheological characterisation, both genotype (p = 0.004) and the rheological protocol (p = 0.042) influenced muscle fibre CSA (**Fig. 9A, E**), with *mdx* muscle fibres remaining larger than WT muscle fibres and a higher compressive force increasing the CSA. These data indicate higher compressive force (40-50% loss in muscle thickness) damages dystrophin-positive and dystrophin-negative skeletal muscle equally, resulting in larger fibre sizes.

### Study 7

This study aimed to investigate the impact of eccentric contraction-induced strength loss on the biomechanical properties of the tibialis anterior. The isometric strength of the anterior crural muscles did not differ between WT and *mdx* mice (**Fig. 10A**; p = 0.209). Because compression at 40% or greater caused muscle damage, we decided to use the non-damaging 0.1 N compressive force to ensure we captured the effect of the exercise-induced injury independent of rheological injury. Fifty eccentric contractions induced 25% and 72% loss in eccentric strength (**Fig. 10B**; p < 0.001) and 32% and 87% loss in isometric strength in WT and *mdx* mice, respectively (**Fig. 10C**; p < 0.001). Eccentric contractions did not affect the thickness of WT or *mdx* muscle (**Fig. 10D**; p = 0.129), but there were compressive force x genotype and compressive force x injury interactions (**Fig. 10E**; p < 0.001), with the injured tibialis anterior of both genotypes compressing less relative to the contralateral uninjured tibialis anterior and *mdx* compressing less than WT. Storage modulus over shear strain was affected by a strain x genotype x injury interaction (**Fig. 10F**; p < 0.001), with storage modulus increased 76% in the injured vs uninjured *mdx* muscle and no change in WT muscle (**Fig. 10G**). Loss modulus was affected by a strain x genotype interaction (**Fig. 10H**; p = 0.005), with injured muscle presenting with a lower and higher loss modulus in WT and *mdx* mice respectively, relative to their uninjured contralateral muscle (**Fig. 10I**; p ≤ 0.004). Loss factor increased with increasing shear strain and was affected by the eccentric contractions (**Fig. 10J**; p = 0.002). When analysed collectively, loss factor was lower in the injured tibialis anterior of both genotypes relative to the uninjured tibialis anterior (**Fig. 10K**; p < 0.001). Eccentric contractions did not affect the change in thickness of the tibialis anterior of both genotypes at the cessation of the protocol (**Fig. 10L**; p = 0.286). These data indicate that eccentric contraction-induced strength loss increases the stiffness of dystrophin-deficient skeletal muscle and its capacity to dissipate energy when deformed.

**Figure 10.**
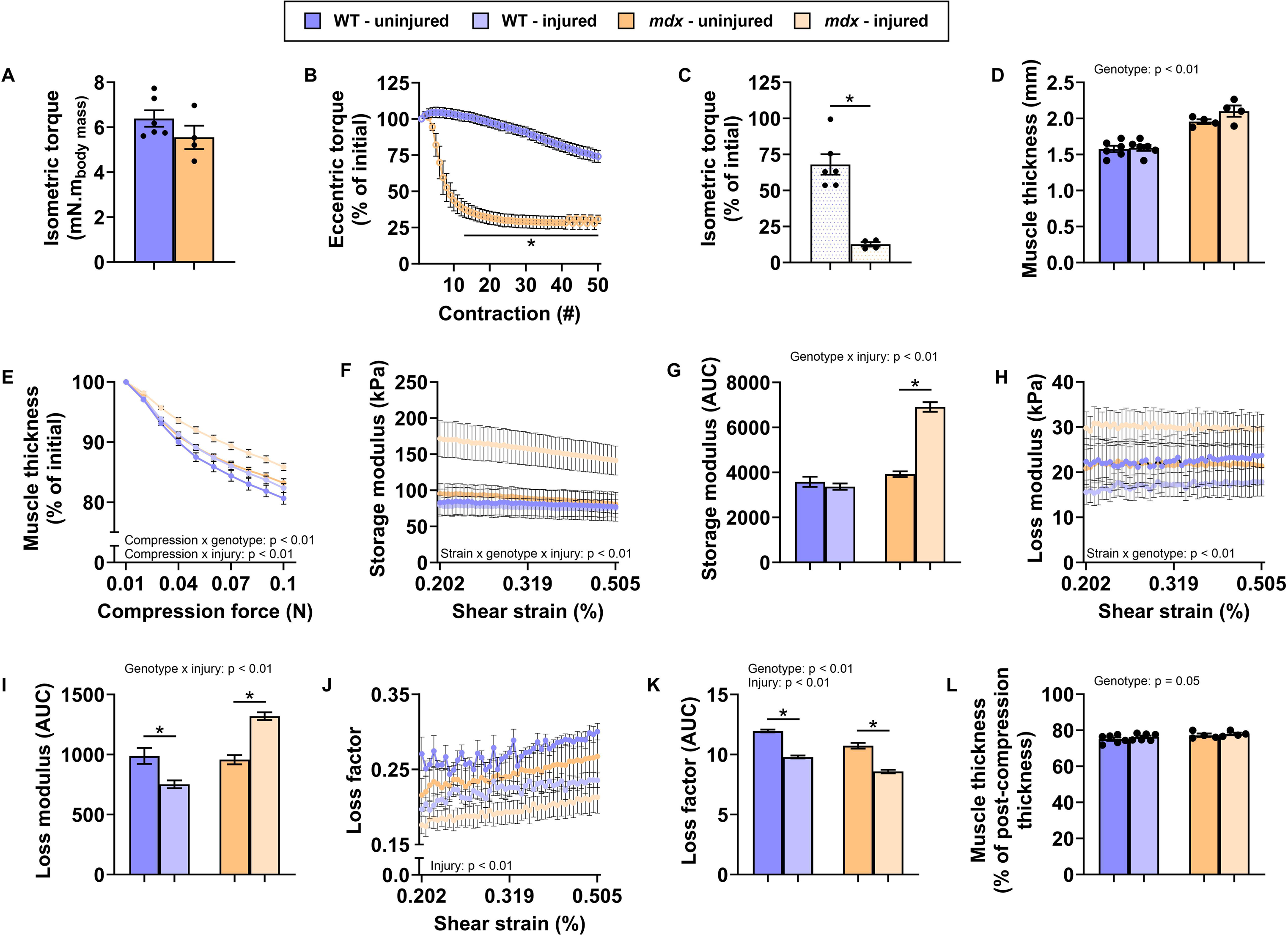
Eccentric contractions stiffen dystrophin-deficient skeletal muscle. A) Isometric strength, B) eccentric strength over repeated contractions and C) isometric strength loss after 50 contractions of the anterior crural muscles of wildtype (WT) and mdx mice. D) Muscle thickness prior to rheological testing, E) muscle thickness as a function of compression force, F) storage modulus over amplitude sweep, G) area under the curve of storage modulus, H) loss modulus over amplitude sweep, I) area under the curve of loss modulus, J) loss factor over amplitude sweep, K) area under the curve of loss factor and L) muscle thickness at the end of the test of injured and uninjured WT and mdx tibialis anterior muscle. Rheological parameters were as follows: compressed with 0.1 N, with amplitude sweeps between 0.001% and 5% at 1 Hz. Data is mean ± SEM. N = 4-6/genotype. * P < 0.05.

## Discussion

The purpose of this study was to examine the impact of dystrophin-deficiency on the *in vivo* murine skeletal muscle biomechanics by assessing muscle viscoelastic properties. The response of live murine dystrophin-positive and dystrophin-negative tibialis anterior to dynamic compressive and rotational strains was able to be evaluated using a purpose-built rheological testing rig, termed “myomechanical profiling”. The developed *in vivo* rheological technique was able to distinguish differences between the dystrophin-positive and dystrophin-negative muscles’ response to applied deformation, stiffness, elasticity, and relaxation, indicating dystrophin-deficiency alters the viscoelastic properties of skeletal muscle.

### Technological innovation

While previous studies have sought to characterise the viscoelastic properties of dystrophin-deficient skeletal muscle, their scope was largely confined to *ex vivo* analyses of excised muscles and investigations at the level of individual muscle fibres or cells [16, 17, 22, 38, 39]. One study has attempted to characterise murine tibialis anterior muscle using rheology *ex vivo*, with tissue samples excised and sectioned for testing [22]. However, the impact of dystrophin deficiency on whole-muscle biomechanics *in vivo* has remained insufficiently explored due to the absence of technologies capable of facilitating live measurements – a critical caveat because components of the *in vivo* environment (e.g., blood supply, physiological temperature and tendon attachments) likely impact biomechanical functionality. Several technical challenges have impeded the development of an effective *in vivo* assessment method. These include the need to accommodate constraints imposed by the murine skeletal system, the isolation of live muscle for controlled load application, and the alignment of muscle fibre and fascicular organization with the directionality and characteristics of mechanical loading. Additionally, ensuring high measurement accuracy, repeatability, and sensitivity is essential for detecting genotypical differences in muscle response. Furthermore, minimising invasiveness and muscle damage during assessment is critical to preserving tissue integrity for subsequent physiological evaluations and potential therapeutic interventions. We have developed a novel approach to enable the *in vivo* rheometric measurements of murine tibialis anterior skeletal muscle. To the best of our knowledge, this is the first study to conduct *in vivo* rheological testing of murine muscle while maintaining blood supply and neuromuscular connectivity. This innovation allows for more physiologically relevant measurements, avoiding tissue excision and preservation artefacts.

The development and final construction of the *in vivo* rheological testing required several critical design considerations: the mouse needed to be held stationary in a supine position to allow for the rheometer probe access, a fixture for an isoflurane nose cone needed to be integrated into the design to ensure continuous anaesthesia during testing, a heated platform needed to be incorporated to maintain normal mouse body temperature and minimize physiological stress, and the apparatus needed to be designed to restrain the mouse without obstructing circulation, preserving the blood flow to the tibialis anterior muscle throughout testing. A key challenge was isolating the live tibialis anterior muscle from surrounding structures such as bones, tendons, and adjacent muscles for testing. To achieve this, a small metal plank was designed to slide underneath the tibialis anterior. This plank was thin with carefully smoothed edges to fit with zero surrounding tissue damage while being sufficiently sturdy to support the muscle during rheometric testing without flex.

The apparatus was designed to be adjustable for inter-mouse anatomical variability, with feed screws allowing fine-tuned positioning in both vertical and horizontal axes for the knee clamps, footrest, and the mouse bed. This ensured that each mouse tibialis anterior could be precisely and reproducibly aligned under the rheometer probe. To further enhance testing consistency, the foot and tibia, as well as the tibia and femur, were positioned at a fixed 90° angle to ensure that the tibialis anterior muscle was maintained at their optimal length. The foot was secured to a footrest on the apparatus with tape, while the knee was held in place by a clamp. The latter ensured the tibia was positioned horizontally and the femur was vertical during testing. This alignment minimized variations in baseline muscle tension, enabling more reliable comparison between mice. The combination of these design elements resulted in a robust, adaptable system for conducting *in vivo* rheometric testing of murine muscle, providing a new methodological framework for studying muscle biomechanics under physiological conditions.

The *in vivo* rheological profiling protocol to quantify the effects of dystrophin-deficiency on murine skeletal muscle was also developed. The optimal rheological testing parameters, including normal force, angular shear amplitude and shear rate were explored. It was key to prevent the probe slippage on the surface of the muscle during testing, while minimising muscle damage from the level of compression and shear applied during testing. The latter was particularly important for the tests involving eccentric contractions, so that any changes observed through the rheological measurements could be attributed to the effect of damaging eccentric contractions alone. **Figure 2** shows strain amplitude sweep tests conducted at 0.01 N, the lowest force the rheometer was able to detect, of normal force at four different frequencies. Probe slippage was apparent at higher frequencies, both visually and from the erratic nature of the data readout. The 1 and 0.1 Hz tests were deemed stable, whereas at 10 Hz and 20 Hz were avoided as the probe slippage was observed. Increasing the force to a moderate 0.1 N improved probe-muscle contact allowing to extend the range of rotational shear amplitude without probe slippage and tissue damage. The latter was confirmed using EBD staining, which showed that 0.1 N of compression up to 5% rotational shear strain did not result in higher levels of muscle damage relative to controls. Furthermore, both WT and *mdx* had similar levels of damage, independent of dystrophin loss, when compression was >40% of muscle thickness.

### Muscle stiffness

Rheological profiling results can be correlated with previous observations that dystrophin-deficiency causes skeletal muscle pseudohypertrophy in the C57BL/10-*mdx* mouse [13], often attributed to the deposition of fat and connective tissue within the muscle. The increased stiffness of *mdx* tibialis anterior compared to WT as observed in both static compression and compression strength testing to a specified compression level can be attributed to the hypertrophic structural changes. Rheological measurements of stiffness using storage and loss modulus confirmed this genotypic disparity - both G’ and G” were consistently higher in *mdx* tibialis anterior muscles relative to WT. The differences in G’ were more pronounced at lower shear strains but narrowed at higher strains, whereas the difference in G” remained relatively stable across all strain levels. G’ exhibited higher shear strain dependence in *mdx* mice, whereas G” did not, within the range of strains tested. This strain dependence of the anterior tibialis elastic component response indicates the underlying hypertrophic structural changes being a dominant factor at the lower shear strains but not contributing as much to the elasticity of the muscle at higher shear strains. However, the viscous component of the muscle response appears to be independent of shear strain, at least in the strain range tested. Our data indicates that the increased stiffness of *mdx* tibialis anterior is associated with its collagen content. This supports the hypothesis that fibrosis contributes to the increased *mdx* muscle stiffness in both compression and shear.

Collagen is a structural protein found in a wide variety of connective tissues, including skin, tendons, ligaments, and the extracellular matrix of muscles. While it plays an important role in maintaining tissue integrity, excess collagen accumulation in muscle tissue, which occurs in DMD [40], leads to fibrosis [8] - the architecture of the latter has been shown to stiffen the muscle [4, 5]. Antifibrotic therapies such as Pamrevlumab are currently under investigation to mitigate collagen buildup in DMD [41–43]. Although preclinical trials in mouse models have shown promise [44–48], clinical efficacy in humans has yet to be confirmed [41–43]. In this study, we measured muscle stiffness using G’ and G” and indirectly assessed collagen levels via hydroxyproline content. Since the precise collagen content within the 3 mm cylindrical muscle mid-belly section could not be isolated, whole muscle hydroxyproline values were reported instead. Hydroxyproline content showed a positive association with G’, and to a slightly lesser extent, with G”. Given the structural role of collagen, its buildup likely contributed to the increased stiffness of the hypertrophic *mdx* muscle tissue, because fibrosis, characterized by enhanced ECM crosslinking, results in architecture within the muscle that resists deformation [49, 50]. Moreover, as the testing was conducted *in vivo*, the measured stiffness is likely influenced by the collagen content throughout the muscle, not just the region directly beneath the rheometer probe. The mechanical impact of collagen is not limited to the immediate contact region between the probe and the muscle due to the strain distribution within the entire muscle, illustrated conceptually in **Figure 11**.

**Figure 11.**
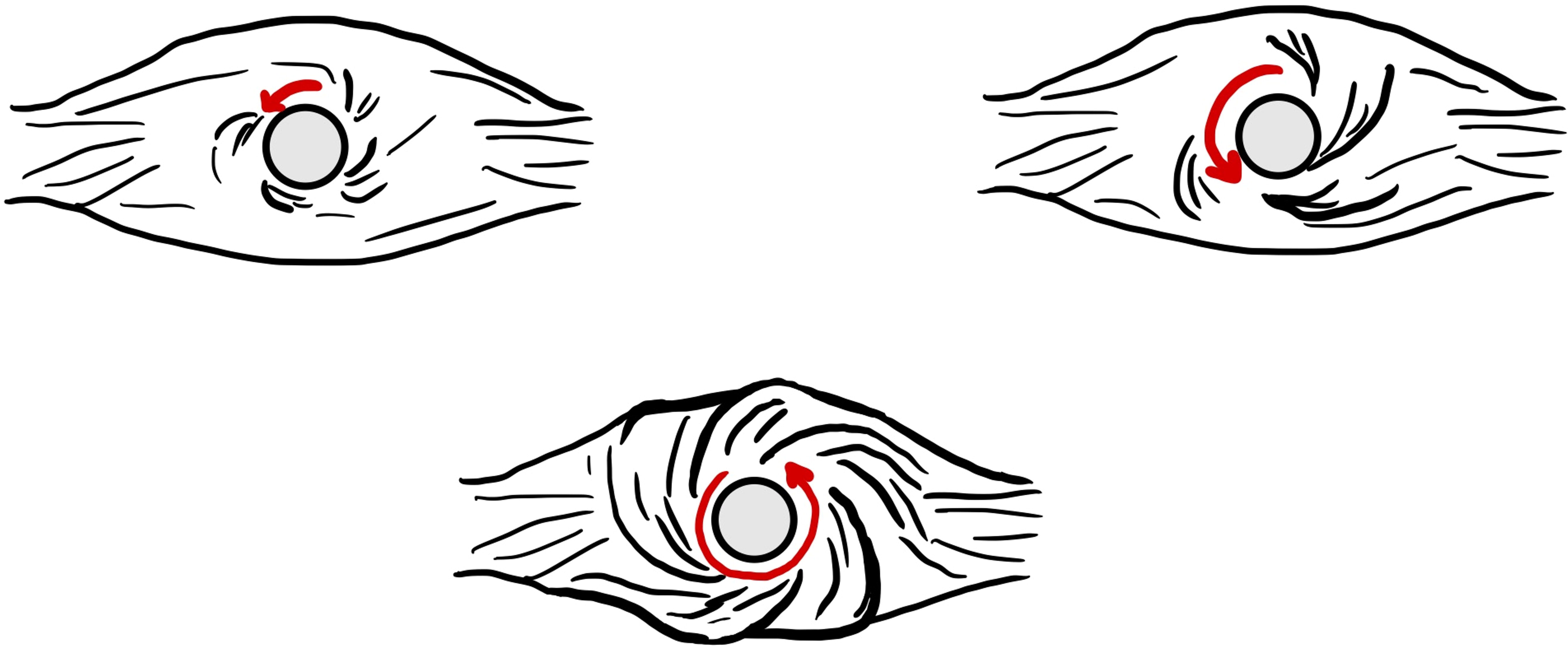
Conceptualised effect illustrating the hypothesised association between strain distribution and the corresponding effects on muscle biomechanics (not to relative to scale between probe and muscle).

### Muscle elasticity

Muscle elasticity was determined by its ability to recover in thickness following compression and rotational shear and we hypothesised that the loss of the shock absorbing properties of dystrophin [6] would render skeletal muscle compromised. When the tibialis anterior was compressed to 40% of its original thickness, WT muscles recovered to about 90% of its original thickness, while *mdx* muscles recovered to only 82%. This suggests that while *mdx* muscle is stiffer than WT and does not deform as easily, the WT muscle has a greater ability to recover from deformation. Study 4 followed a similar protocol to Study 3, but with an increased shear strain reaching up to 10% instead of 5%, while the level of compression was maintained at 40%. This reduced the difference between genotypes, with both recovering to approximately 80% of their original thickness. This suggests that at higher shear strains, WT muscles elasticity decreases, while *mdx* muscles show minimal change. In Study 5, when repeated 5% shear strain was applied, *mdx* and WT muscles recovered similarly, with *mdx* showing a more elastic response. Interestingly, Study 6 showed that *mdx* muscles had a higher elasticity than WT following a compression to 50% over 30 minutes, and that under this strain maintenance, there was no difference in the relaxation behaviour between genotypes, suggesting the capacity of muscle to displace under constant load is independent of dystrophin deficiency or the dystrophic environment. These results suggest that under rotational shear strain, *mdx* muscle are less capable of completely recovering after deformation, whereas in pure compression, it recovers comparably or better than WT muscle. This difference may be linked to the internal collagen architecture and how it responds to different modalities of mechanical loading, which requires further exploration.

### Contraction-induced damage and rheological profile

Dystrophin-deficient *mdx* vertebrates exhibit pronounced strength loss and muscle fibre damage following eccentric contractions [5, 32, 33, 51, 52]. Here, we show that an eccentric contraction protocol inducing 85% strength loss in *mdx* muscle further stiffens *mdx* muscle. Eccentric contractions had no meaningful effect on G’ in WT muscle, which is expected given the minimal strength loss observed in this genotype. In contrast, G’ increased markedly in *mdx* muscle post-injury. However, G” decreased in WT after eccentric contractions but increased in *mdx*. The change in G’ and G” following eccentric contractions in *mdx* muscle likely reflects the multi-factorial mechanism of strength loss, which includes mechanical disruption to the muscle architecture [32]. Interestingly, increased G” in *mdx* suggests that muscle damage and, likely, the associated inflammatory response contribute to the muscle’s capacity to dissipate energy. In WT, the reduction in G” suggests decreased energy dissipation properties. These findings show that rheology captures not the structural differences from dystrophin loss but rather the physiological state (quality) of the muscle following contraction-induced strength loss and its mechanical consequences. This suggests that rheological measurements may be able to indicate muscle health, providing insight into damage, repair, and biomechanical function *in vivo*.

## Conclusion

This study developed a novel platform to measure mechanical properties of murine muscle tissue *in vivo*. Dystrophin-deficient *mdx* tibialis anterior muscle was found to be consistently stiffer than WT, which was associated with collagen content. This study shows rheology’s ability to assess the ‘quality’ of skeletal muscle in health and disease.

## Limitations

While the testing was done *in vivo*, the skin of the mouse was removed for testing, something that will affect the local physiology. However, this was necessary to isolate the tibialis anterior. The skin removal would likely cause drying, which may affect some of the measurements. The tibialis anterior was the only muscle tested, though this was justified as it is routinely tested for *in vivo* physiology of the *mdx* mouse, is easily accessible and can be isolated. Lastly, while this study did measure collagen content using hydroxyproline as a proxy, this is purely a quantitative assessment, and we did not investigate the specific architecture of the collagen within the muscle. Future research could investigate this relationship to draw stronger conclusions regarding collagen’s role in affecting G’ and G”.

## Acknowledgements

The authors would like to thank Dr Ashish Kumar, Helena Barnes and Marzieh Monfared (Anton-Paar Australia and New Zealand) for the loan, installation and technical advice associated with the rheological characterisation of the tibialis anterior muscle using the MCR702e (Anton-Paar). The authors also acknowledge Lauren Hunt’s (University of Canterbury) contribution to the mouse apparatus prototype development. The authors would like to thank Alice Cerdeira (University of Canterbury) for sketching the illustrations used in the graphical abstract and Figure 11.

## Author contributions

A.L. and N.K. conceptualised the study. All authors performed experiments. P.D., N.K., and A.L. analysed and interpreted data. P.D., N.K., and A.L. prepared the manuscript. All authors read and approved the final manuscript and agree to be accountable for all aspects of the work in ensuring that questions related to the accuracy or integrity of any part of the work are appropriately investigated and resolved. All persons designated as authors qualify for authorship, and all those who qualify for authorship are listed.

## Data availability

Data will be made available upon reasonable request of the corresponding author.

